# Learning to see surface properties by predicting visual experience

**DOI:** 10.64898/2026.11.20.768077

**Authors:** Fatma Kiliç, Roland W. Fleming, Katherine R. Storrs

## Abstract

The central challenge of perception is inferring the structure of the outside world from ambiguous sensory information. Here we find that an unsupervised neural network trained solely to predict forthcoming sensory signals spontaneously learns to untangle various physical properties, including surface reflectance and various kinds of object motion. Individual units in the model developed sensitivity to specific static and dynamic image cues that are diagnostic of surface properties, such as how highlights slide and change shape as an object rotates. We found that linearly decoding reflectance from the model’s latent space reproduced human reflectance judgments, including striking illusions that occur when reflected patterns are rigidly ‘stuck’ to the surface, leading humans and the model to interpret them as matte texture patterns. Taken together, our findings suggest that the visual system could learn to infer distal world properties entirely through passive observation, just by predicting what sensory signals come next.

## Introduction

A fundamental challenge for the visual brain is working out the structure of the outside world from the continually transforming patterns of sensory stimulation that arrive on the retina. In order to act on the world, the visual system must deliver information about the size, shape, layout, and material properties of surfaces and objects in the surroundings. Yet the raw sensory signals are highly ambiguous about their distal origins in the world.

Consider the challenge of determining what material an object is made of. An object like the one in **Figure 1**, with a white streak across the surface, is consistent with at least two distinct underlying causes. The feature could be due either to texture (e.g., a pattern painted on the surface) or to the object reflecting its surrounding environment. Not only is the correct solution not directly observable—the two possibilities can give rise to identical images—but the brain must also learn that these two distinct causes of surface markings exist in the first place: texture and reflection.

**Figure 1:**
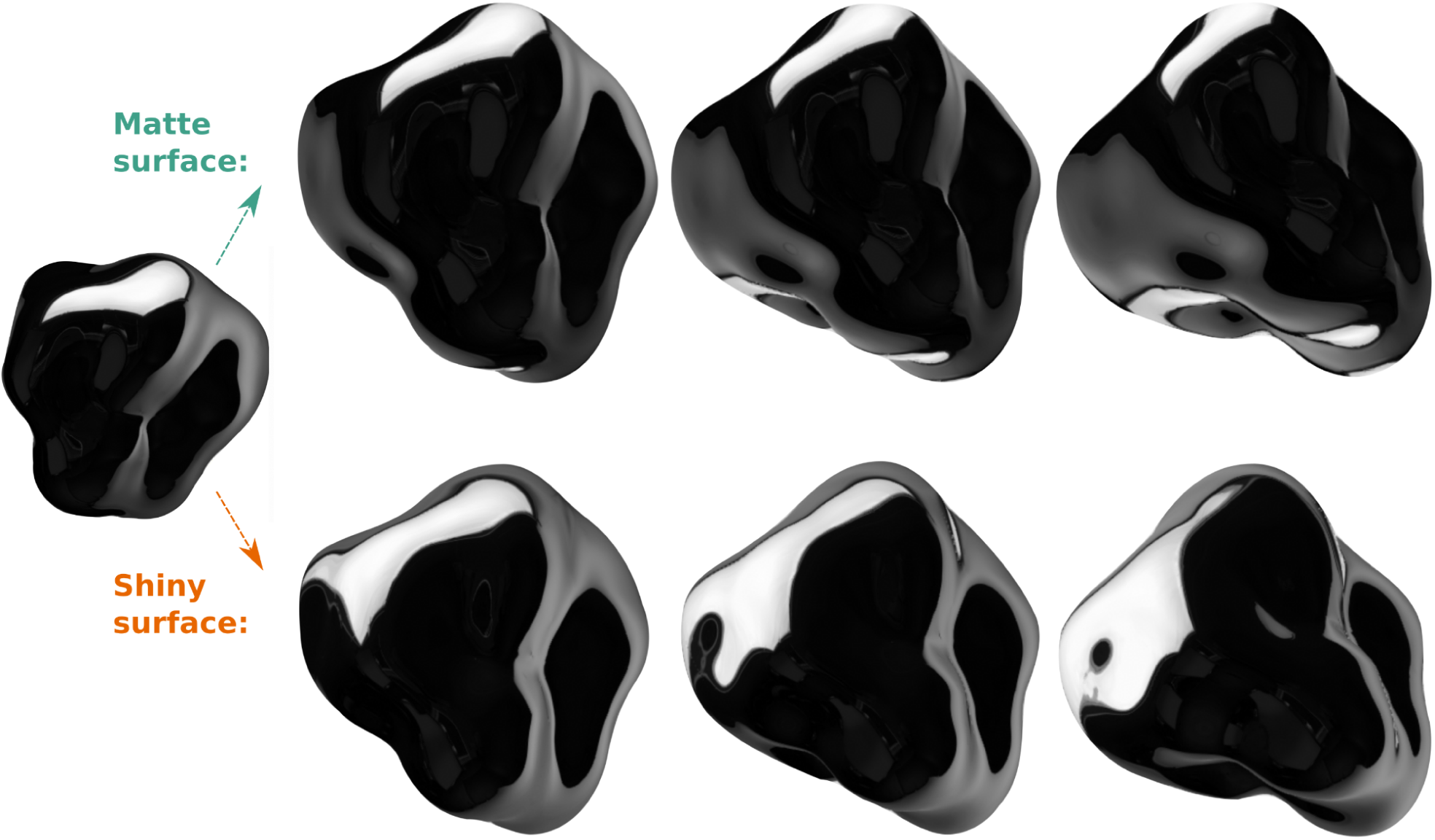
Making inferences about the causes of sensory experience. The material properties of the object on the left are ambiguous. The white streak could be either a texture marking “painted onto” the surface, or a reflected highlight. One needs to know which of these is the case in order to predict how the object will look when it moves. A texture marking on a matte surface (top row) will stay “stuck” to the object as it rotates. A reflection on a shiny surface (bottom row) will change shape and size as it slides across the object. Our modelling approach reverses this logic: by learning to predict moving objects, does a system learn the existence of matte and reflective materials?

We propose that brains learn about the existence of different types of distal material by learning to predict how proximal visual features vary over time and space. We previously showed that, in the static domain, perception and misperception of surface reflectance are well predicted by representations within an unsupervised feedforward deep neural network that has learned a model of the spatial structure in images (Storrs et al., 2021). Temporal prediction offers a wealth of additional visual structure from which to discover and untangle scene properties. Specular surfaces reflect light at a specific angle proportional to the angle of incidence and tend to appear shiny or mirror-like, whereas surfaces that scatter light diffusely tend to appear matte. When an object moves, the reflections on a specular surface rush across its surface, locally growing or shrinking as they encounter convexities and concavities in the object’s shape (Adato et al., 2010; Doerschner et al., 2011; Fleming et al., 2004; Koenderink & van Doorn, 1980). In contrast, matte surfaces create more homogeneous patterns of motion, for example, tracking the steady displacement of texture markings in the direction of object rotation. Therefore, to make good predictions about future proximal images, a system would benefit from accurately modelling the distal causes such as surface reflectance. To reverse this logic, predicting the way image features such as highlights and textures change over time becomes an objective by which the visual system could learn the existence of different types of material in the first place, without receiving explicit labels.

In the fully-developed visual system, motion is indeed a powerful cue to material properties. If asked to evaluate the reflectance of objects, observers typically move them back and forth to observe the way highlights shift and change shape across the surface (Lin et al., 2025). We suggest that the brain not only uses motion cues to recognise materials, but that predicting these motion signals is how it learns, in the first place, the existence of material categories that cannot be learned from spatial relationships alone. Here, we therefore tested whether learning by temporal prediction in a recurrent network discovers the motion cues that humans also rely on to judge surfaces. The key hypothesis is that end-to-end learning of the statistical structure of motion patterns can discover—in an entirely unsupervised way—the core ontological distinction between matte and glossy surfaces, by mapping stimuli with different reflectances to distinct locations within the model’s latent space (Bengio et al., 2013; DiCarlo & Cox, 2007; Lotter et al., 2017; Storrs et al., 2021). Moreover, if humans also learn the key cues in a similar way, we would expect such a model to reproduce the kinds of judgments humans make when presented with moving patterns.

To test this, we leveraged advances in recurrent neural networks to create an unsupervised model that learned visual representations from unlabelled videos by predicting how objects change in appearance as they move. By rendering a dataset of artificial videos for which ground-truth scene properties were known, we were able to fully control the visual diet of the network, and subsequently probe its ability to untangle those scene properties. Although pixel-wise prediction may seem like a low-level learning objective, there are intuitive reasons to think that it may drive a system to learn far deeper visual representations. Specifically, in order to predict what happens next, it is very helpful to understand the scene in terms of higher-level causes—objects with certain geometries, textures, and materials, starting in certain poses, and moving along certain trajectories.

## Results

We rendered 20,000 twenty-frame videos of simple procedurally-generated scenes in the Unity3D graphics engine. Each scene consisted of one of 10 objects, steadily translating and rotating with random direction and velocity in front of a uniform grey background of random albedo (**Figure 2**). Objects were illuminated by one of 10 real-world light fields and had either a mirror-like surface material or a matte surface textured with one of 10 patterns (**Figure 2**). Objects were positioned close to the virtual camera at a slightly jittered distance and a random pose, and were rotated by a random angle up to 180 degrees along all three axes over the course of 20 frames. This created video clips of close-up, smoothly curved moving surfaces with either matte or specular optic flow.

**Figure 2:**
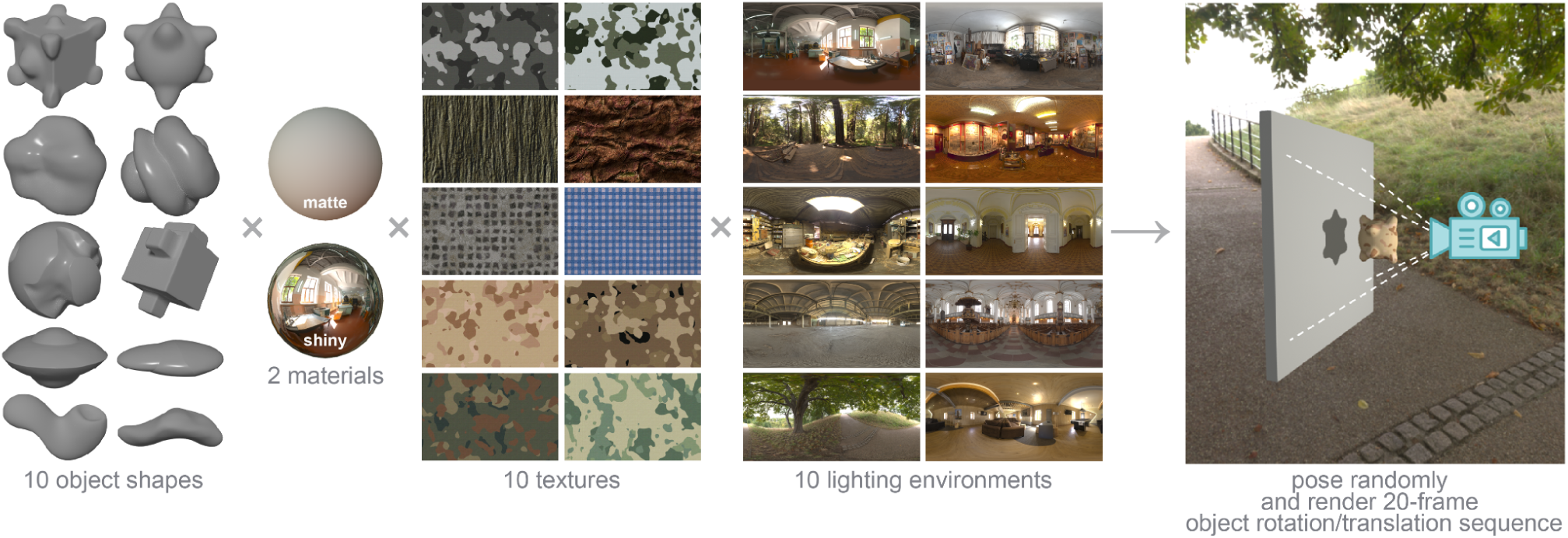
Generative world for rendering videos of moving objects. From left to right: To create each video, a random object geometry, with either a mirror-like surface material or a random texture pattern, was placed in a random illumination field and filmed in front of a blank wall rotating or translating along a random trajectory for 20 frames (see Training Dataset in Supplementary Videos).

### Models learn to predict the complex behaviours of specular highlights

We trained a four-layer recurrent network to predict the pixels of the next frame in each video. Specifically, we trained a PredRNN (Wang et al., 2017) recurrent convolutional network (**Figure 3a**) to minimise mean squared pixel error between predicted and actual future frames. For frames 1-10, the network predicted one frame into the future; after frame 10, it predicted until frame 20 with no further input.

**Figure 3:**
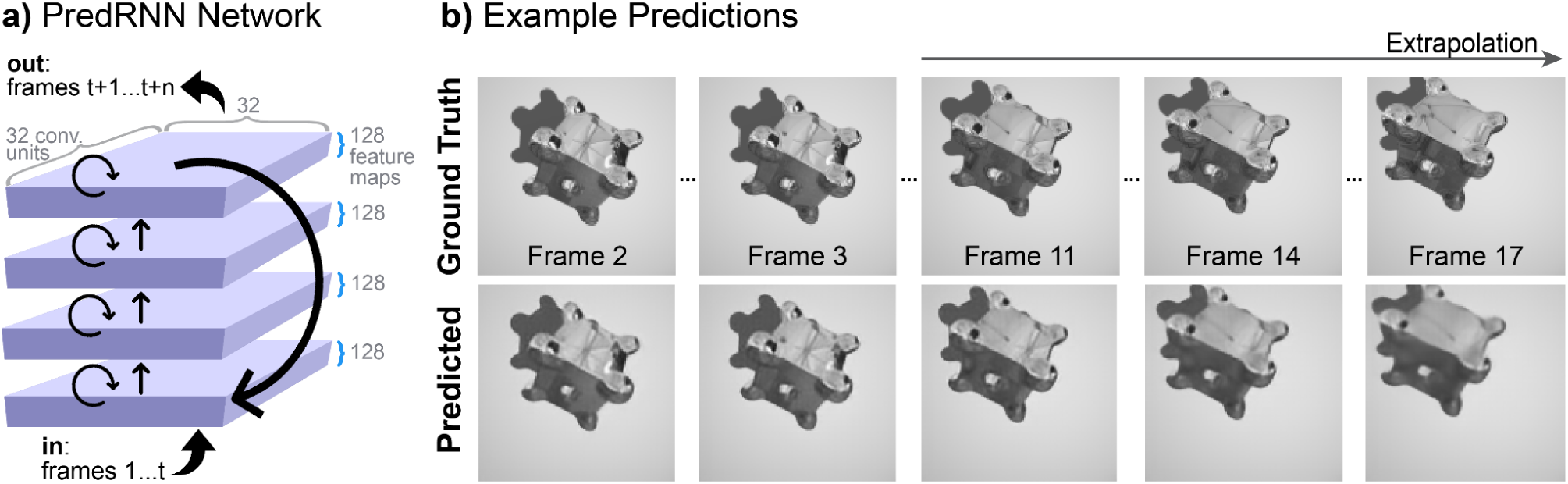
PredRNN video prediction network. **(a)** The architecture consists of four layers of convolutional feature maps containing recurrent LSTM units. The input and output layers are of the same dimensionality as the video frames. The training objective of the network is to minimise the mean squared pixel error between the predicted next frame(s) and the true next frame(s). **(b)** Selected frames from an example sequence of ground-truth (top) and network predictions (bottom) for a video not seen during the network’s training.

The trained PredRNN was exceptionally good at predicting future frames of novel videos featuring previously unseen combinations of object, lighting, material, and motion trajectories. The network’s average MSE was 0.003 per frame, compared to 0.018 for the persistence baseline of simply taking the previously predicted frame as the prediction of the next (starting from the last observed frame), and 0.008 for an optical flow-based baseline on RAFT (Teed & Deng, 2020). The network could extrapolate up to ten frames beyond its input data with good quality (**Figure 3b**).

Studying individual frames reveals that the trained PredRNN model was able to anticipate highly nonlinear changes between frames, such as the rapid expansion or contraction of specular highlights as they slide across a moving object’s surface. Extrapolated frames correctly captured the different behaviours of the two distinct reflectance conditions.

### Models learn an untangled representation of underlying scene properties

Our primary interest was whether the low-level objective of pixel prediction would drive the unsupervised model to discover underlying scene properties, since these are the generative factors that give rise to the moving image data. Exploring the organisation of internal representations via t-SNE, we found that over time, scenes clustered according to whether objects translated or rotated (**Figure 4a**). The representation of rotating objects further revealed separate clustering for matte and shiny materials, whereas the representation of material was intermixed for translating and static objects. This pattern is consistent with the fact that matte and specular optic flow patterns differ most for rotating surfaces, which cause specular reflections to dramatically warp and slide.

**Figure 4:**
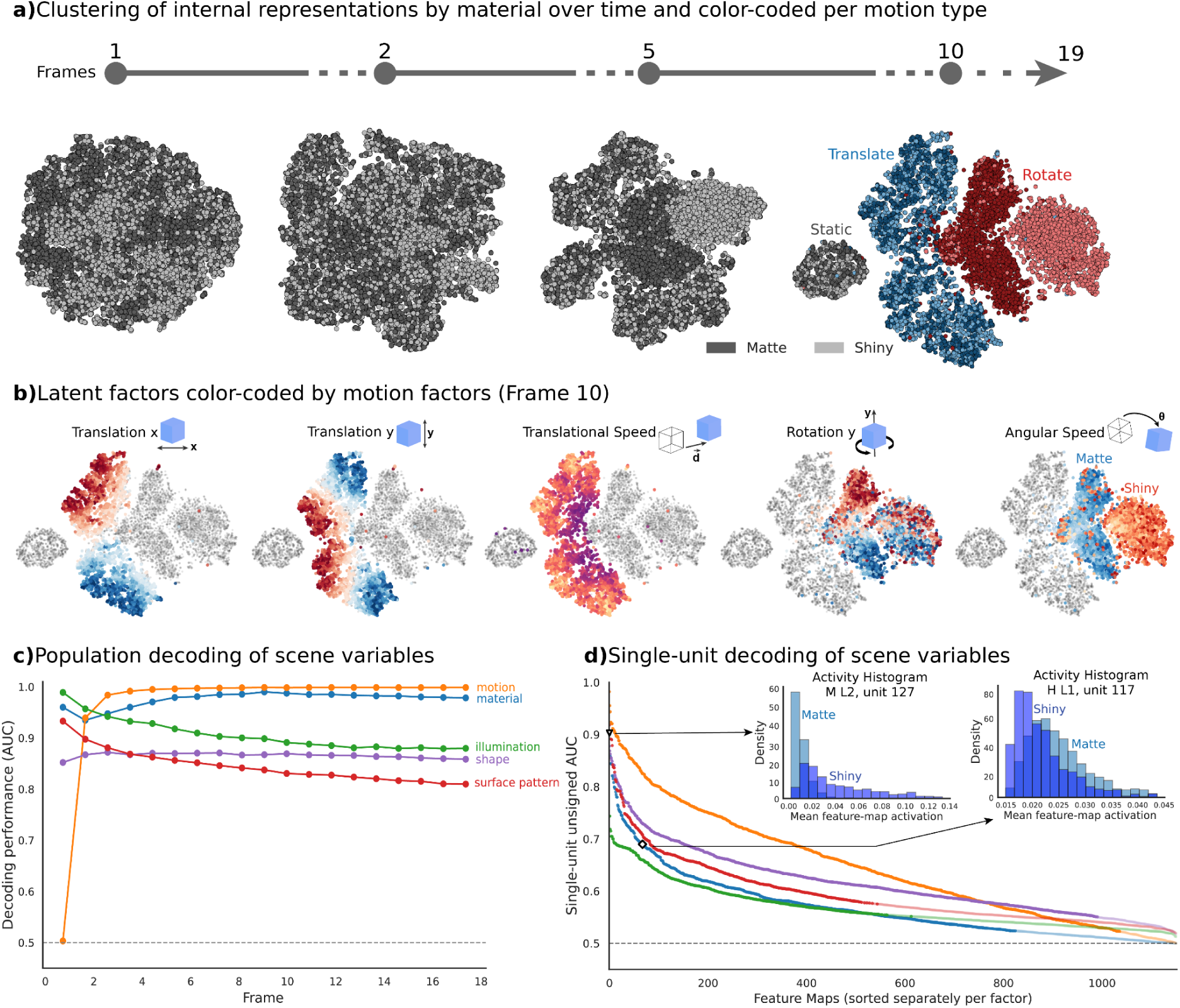
Internal PredRNN representation of material- and motion-related structures. **(a)** 2D tSNE embedding of 9500 scenes in layer 2 of the PredRNN network across selected time frames. Early grayscale frames show the gradual emergence of material-related structure. Grey value indicates whether the scene contained a matte (darker) or mirror (lighter) surface. In Frame 10, additional colour indicates the motion type: translating, rotating, or static; **(b)** Frame 10 embeddings of internal representations of the network coloured by continuous motion factors: translation along the x- and y-axes, translational speed, rotation around the y-axis, and angular speed. Gray points indicate the scenes for which that motion factor is not applicable. For translation and rotation, colour represents the direction. For translational speed, the colour intensity indicates the magnitude of total translational displacement during the video (purple = slow translational speed and yellow = fast translational speed). For angular speed, the colour (blue or orange) indicates the material, while colour saturation shows the magnitude of total angular displacement; **(c)** Linear decoding analysis across frames, quantified by the area under the receiver operating characteristic curve (AUC) per scene variable. Dashed line indicates the chance level; **(d)** Single-unit decoding of each unit (feature map), defined as the directionless single-unit AUC, across all layers and memory types for each scene property. Units are sorted independently for each factor. The dashed line indicates the chance level. Dark circles indicate units with statistically significant AUC values (permutation test, FDR-corrected q ≤ 0.05), whereas light circles indicate units without significant decodability. The histograms represent the activity distributions of two example feature maps with opposite material preferences.

The network revealed fine-grained encoding of motion direction and speed nested within the broad clusters of motion type and material. For translating objects, translation along the x- and y-axes and overall speed were smoothly organised within the network’s representational space (**Figure 4b**). For rotating objects, magnitude of out-of-plane rotation was smoothly organised across both matte and shiny objects, and angular speed formed smooth continua within the separate clusters of matte and shiny materials. This is consistent with angular speed having distinct visual consequences depending on the object’s material, as rotation creates different optic flow patterns for matte and specular surfaces. Overall, these findings reveal that the latent space of the trained network forms a highly structured representation of objects according to their kinematic properties, including the interaction between movement and surface material.

To complement the visualization of representational organization, we measured to what degree scene information was available via a linear readout of the network’s internal activations at each time point. Motion, material, shape, illumination, and surface pattern could all be decoded simultaneously from the network’s internal representations substantially above chance (**Figure 4c**). Decoding performance for motion and material peaked only after temporal information from multiple frames had been integrated into the network’s activation state. Other scene variables such as illumination and surface pattern were already maximally decodable at the first frame, and decoding performance slightly declined after several frames. Information about scene variables was present not only at the population level but also at the single-feature level. We evaluated the decoding performance for each feature map of each memory type in each of the last three layers of the network (1,152 features in total; **Figure 4d**). The lowest single-unit decoding performance was for surface pattern, which could be significantly decoded from 44.1% of feature maps, and the strongest was motion, which could be decoded from 88.7%. Importantly, we observed that high decoding performance from different units was driven by preferences for different categories within a scene variable. The histograms in **Figure 4d** illustrate this phenomenon for material: one feature showed higher activation for matte objects, whereas the other showed stronger activation for shiny objects. This suggests that single-unit decoding is not driven solely by greater contrast or motion energy within some categories driving higher responses across all units, but by learned selectivity. Taken together, these results suggest that the information is represented in a temporally dynamic and non-uniform manner, with different variables showing distinctive patterns over time and across feature maps.

### Individual “neurons” develop selectivity for distal properties

Next, we performed feature map ablation analyses to causally understand the role of individual feature maps in the network’s overall scene variable representations. If lesioning a small number of units destroys population-level decoding, it would suggest sparse functional specialisation for that scene property; if decoding is unaffected, it would suggest a fully distributed representation. We found an intermediate result between these two extremes (**Figure 5b**). When we lesion the top 10% of feature maps that carry the most information about that scene variable (i.e., targeted lesioning of the units highlighted in **Figure 5a**), the decoding performance of the network is reduced significantly compared with lesioning randomly selected feature maps (all five scene variables, one-sided empirical randomization tests, Holm-corrected *p* = 0.010). The most pronounced performance drops were observed for material, and the impact was selective — lesioning these units had little effect on the decoding of other properties, and lesioning units that carried information about other variables had less of an effect on material decoding (**Figure 5b**). This suggests that the representation of material is concentrated within a relatively small subset of feature maps, and that these feature maps do not also carry critical information about other properties. Other scene variables appeared to have representations that were less selective, but still distinct. Motion decoding was reduced only by lesioning the 10% of units that carried the most information about motion — but lesioning these also damaged the decoding of other factors. Lesioning shape-informative units selectively impaired shape decoding — but shape decoding was also impaired by lesioning units informative about any other factor, consistent with overlapping representations across different scene factors. Lesioning units important for surface pattern and illumination had less selective impacts on the decodability of these variables, which were also affected by lesioning other units. A direct analysis of the overlap found that nine out of the ten possible scene factor pairs had significant overlap between their most-informative top 10% feature maps (16.4–29.3% overlap; Holm-corrected *p* < 0.05). Overall, these analyses highlight that small subsets of feature maps within the network can contain functionally relevant information regarding a specific scene variable, but that the representation is not fully disentangled.

**Figure 5:**
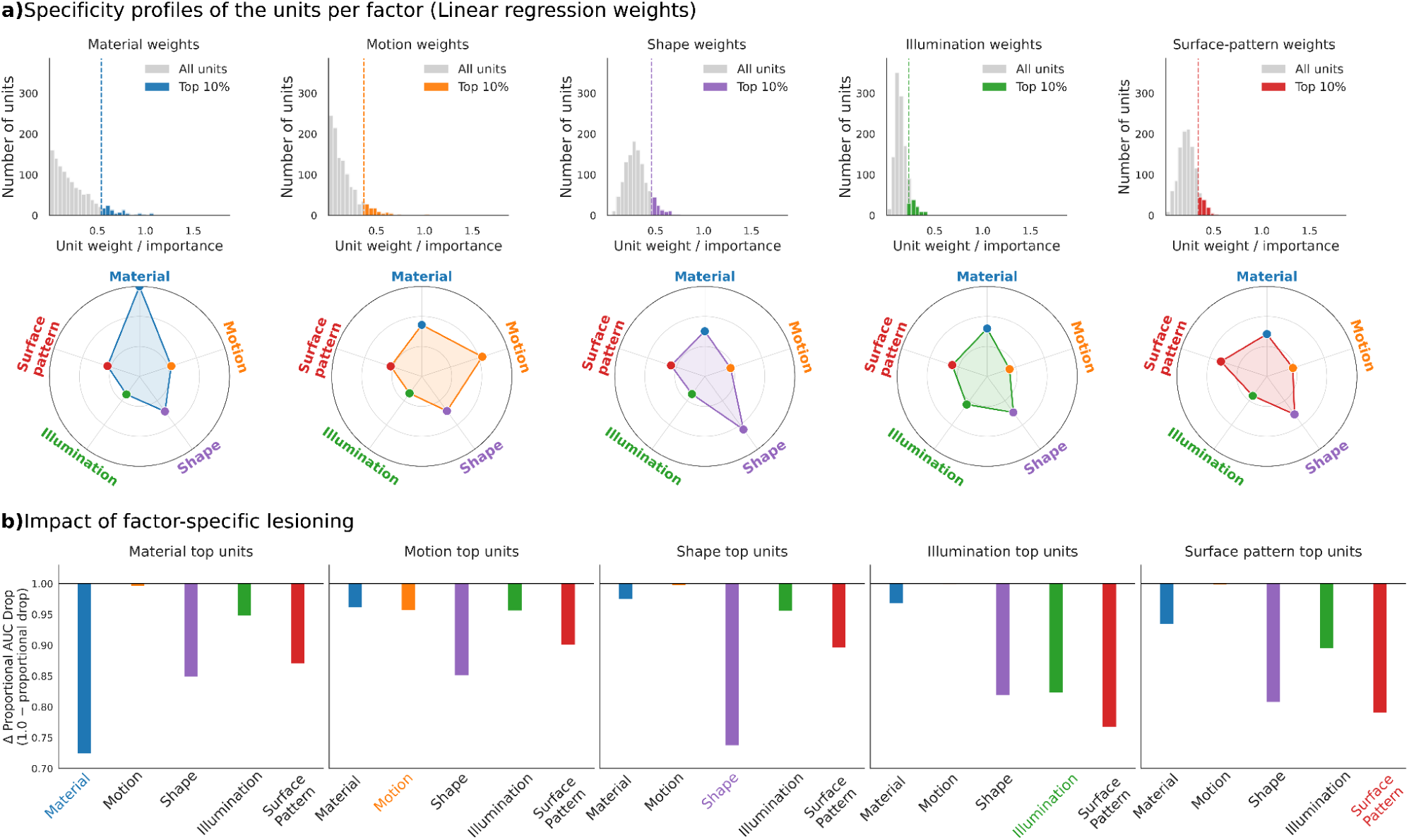
Specificity of units for particular scene properties, and impact of lesioning. **(a)** Specificity profiles across units for each factor, quantified by the weights of a linear decoder fitted to predict each factor. Histograms in the top panel show the distribution of linear decoder weights across all units in the network, for decoders fitted to predict each factor. The coloured bars highlight the top 10% of unit weights. The polar plots below show how factor-specific these units are, by plotting the average weighting those top-10% of units received in each of the other factors’ fitted decoders. For example, the leftmost plot shows that units which received top-10% weights in the material decoder received relatively low weights in the other factors’ decoders. **(b)** Impact of factor-specific lesioning, defined as lesioning the units with the top 10% of weights in the linear decoder fitted to that factor (i.e. those with coloured bars in the histogram above). The impact of lesioning is quantified by the difference in AUC for targeted versus random lesioning.

### Models learn both static and motion cues to material, like humans

Both static and motion cues contribute to material perception in humans. Motion is particularly important to disambiguate different reflectance properties (Doerschner et al., 2011; Hartung & Kersten, 2002; Kam et al., 2015; Marlow & Anderson, 2016). For example, the image in **Figure 1** is consistent either with a shiny object reflecting its surroundings, or with a matte object onto which a complex pattern has been “painted.” While they cannot be distinguished from a single image, motion reveals systematic differences in the kinds of optic flow produced, as reflections or surface patterns move relative to the surface and can resolve these ambiguities. To understand whether the network uses similar temporal information to resolve ambiguities, we created 90 matched sets of stimuli depicting the same object, illumination, and pose, but in which reflections either moved normally (shiny appearance) or stayed “stuck” to the surface (matte appearance), or in which the object remained stationary (Doerschner et al., 2011; Kam et al., 2015). All had identical static material cues, but differed in motion cues. These stimuli allowed us to tease apart the contributions of static and motion cues to material decoding in the model (see Diagnostic Dataset in Supplementary Videos).

We applied the linear decoders trained in the previous analyses to read out material from the network’s internal representation at each time point. Initially, when only the static image was present, all three conditions were represented similarly, and the network could not differentiate between conditions. Yet, as soon as the temporal structure emerged, the network classified the scenes containing objects with a shiny surface as shiny/mirror-like, and those manifesting a static appearance or a “sticky” reflection were classified as matte (**Figure 6a, left)**. This indicates that the network does not solely depend on static visual cues, but benefits from the interaction between surface properties and motion cues.

**Figure 6:**
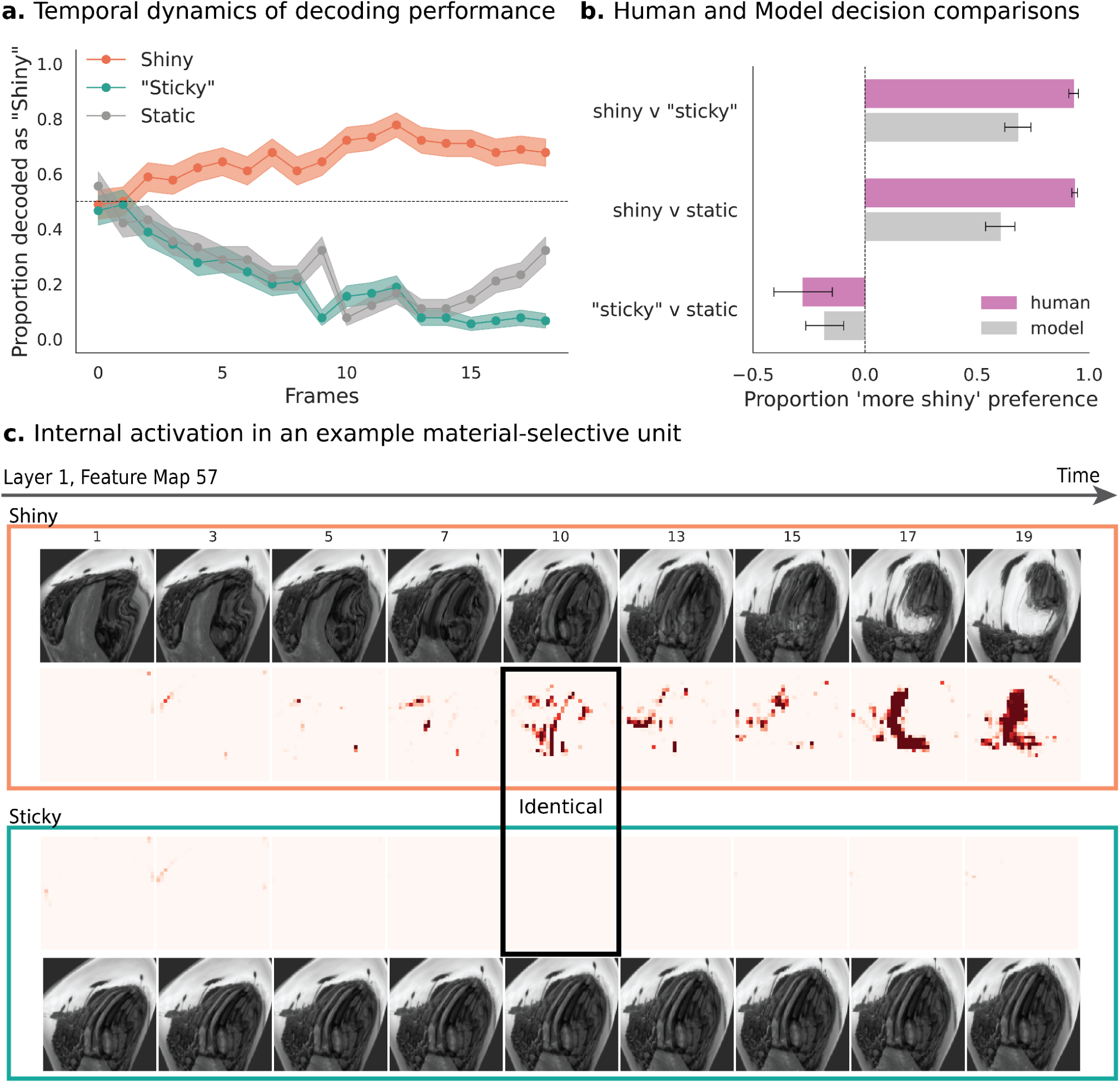
Manipulating material-from-motion cues reveals sensitivity to both static and motion cues to material. **(a)** Temporal dynamics of decoding performance, quantified as the proportion of frames decoded as “shiny” by the decoder. Values above 0.5 indicate that the decoder labels the object in the specified frame as shiny, and values below 0.5 indicate that the object is labeled as matte. **(b)** Relative glossiness (reflectance) decoded from the model agreed with human judgements (N=16) comparing static, moving shiny, and moving “sticky reflection” stimuli; **(c)** Example ground truth and predicted frames for “shiny” and “sticky” versions of one stimulus. Frame 10 is identical in both conditions. The model extrapolates past frame 10 without further input and correctly predicts that reflection markings will slide across (top) or stay fixed to the surface (bottom). Note that, by design, each frame is ambiguous about material, so the reader may not easily be able to distinguish “shiny” from “sticky”. Indeed, frame 10 is identical. Only its past history disambiguates them.

Further, to test whether the network and human observers behave in a similar manner, we collected glossiness/reflectiveness judgments for eight sets of the same stimuli from human participants (N=16). We asked them to compare how glossy/reflective each of the three conditions appeared for each stimulus. The results indicated that human observers agreed with the relative model-predicted reflectance (**Figure 6b, right**), indicating that the manipulation affects the material appearance judgments comparably.

The influence of temporal information on judgments was also evident at the individual feature map level. We compared the network’s activation to the 10th frame of the sequences, which was designed to be identical across conditions. Despite the identical input pixels at frame 10, we found different patterns of activation in the network depending on whether the frame was preceded by a sequence with specular optic flow or matte optic flow. For example, the feature map visualised in **Figure 6c**, which was previously determined by normalised differences in activation strength to prefer shiny surfaces, has a robust response to the frame at the centre of the pictured sequence when it is encountered in the shiny-reflection condition (top row) but not when the same image is encountered in the “sticky-reflection” condition (bottom row). Taken together, the results suggest that the network integrates temporal information with static surface cues, resulting in a similar behavioural pattern to human judgments.

## Discussion

We set out to test whether an unsupervised video prediction network can learn a representation of material that is untangled from object geometry, motion trajectory, and illumination field, simply by learning to predict incoming sensory data. We reasoned that such a network would provide an existence proof that mere passive observation, combined with prediction-based learning, is sufficient to reconstruct (at least a simple) ontology of the outside world. We found that after training on synthetic videos of objects undergoing different kinds of motion, the network successfully separated the distal world properties, such as material, motion, shape, and lighting, based on the proximal sensory input. This occurred without any explicit labels to specify that these were key ontological features of the world it was observing. These findings suggest that spatiotemporal prediction is a sufficiently rich and challenging learning objective to encourage the discovery of world structures that are densely entangled in the raw sensory signals. As well as representing these distinctions in its population response, the network contained individual units that are informative for specific world properties, for example, yielding antagonistic responses for matte vs shiny objects.

Perhaps most intriguingly of all, we also found that the networks reproduced the patterns of shiny vs matte judgments that humans make when presented with stimuli in which we carefully isolate the static and motion cues to surface reflectance. Specifically, we created “sticky reflection” stimuli (Doerschner et al., 2011; Hartung & Kersten, 2002; Muryy et al., 2013) in which reflected patterns were rigidly projected onto the surface of the object, so that they move with the surface—like a texture marking—rather than sliding over the surface the way specular reflections do. Although such stimuli appear shiny in any given frame, when set in motion, they appear matte to human observers. Because such stimuli essentially never occur in the real world, they allow us to tap into the key motion cues that humans use to judge reflectance, independently of *a priori* assumptions based on the spatial characteristics of the image. We found that the unsupervised models also ‘see’ such stimuli as matte, in the sense that they projected to regions of the model’s latent space associated with matte materials, rather than shiny ones. Because humans and the unsupervised models make similar judgments about these stimuli, which were outside their training sets, it suggests that humans might also learn to perceive reflectance through temporal prediction.

There is a long history of previous research on prediction as a central process in perception and neural computation more broadly (Clark, 2013; Friston, 2010; Lotter et al., 2017; Mumford, 1992; Rao & Ballard, 1999; Schultz et al., 1997). The predictive coding framework tends to focus on internal generative models, often describing perception as inference from prediction and sensory evidence (‘perception as controlled hallucination’) (Seth, 2021). Instead, here, we focus on the power of prediction as a learning objective to establish untangled representations, irrespective of how the information is subsequently read out of the latent space (e.g., by a simple discriminative model). In particular, this study shows how prediction-based learning can create and hone the generative model in the first place, and how such unsupervised learning objectives can act as general-purpose mechanisms for uncovering distal ontological structure (‘out there in the world’) from the statistics of proximal sensory signals (Denton & Birodkar, 2017; Grathwohl & Wilson, 2016; Hsu et al., 2017).

We can assess the extent to which information about the outside world is encoded in the network’s latent representation through decoding analyses. Our results revealed that physical scene factors were linearly decodable. Importantly, this information need not be represented by any individual units, but may instead be distributed across the population, although some units do show preferences for certain properties. This is consistent with findings showing decoding of perceptual gloss parameters independently of shape and illumination from cell populations in monkey inferior temporal cortex (Nishio et al., 2012, 2014a). The representation should, therefore, be considered untangled (DiCarlo & Cox, 2007), rather than fully disentangled into unique axes (Nishio et al., 2014b).

Unlike supervised learning objectives, which lead to the emergence of high-level ‘semantic’ gloss and texture units (Morimoto et al., 2026; Prokott et al., 2021; Tamura et al., 2022), we focused here on the bottom-up construction of useful features from low-level image measurements. Previous studies have noted that mirrored objects yield optical flow patterns that are unusual in several key ways, such as the preponderance of optical flow divergence and unmatchable features, larger velocity gradients, and incompatibility with rigid motion (Doerschner et al., 2011). For a network seeking to predict the future trajectory of image patches, such distinctions between reflection-typical and texture-typical optical flow lead to a host of low-level motion cues that they might latch onto. This may help explain the emergence of reflection-selective and texture-selective units in the model, as having acquired sensitivity to a number of easy-to-measure—and mutually correlated—optical flow features.

Our study has a number of limitations. The model operated in the deliberately simplified sandbox of an idealised visual environment. Its world consisted of only a small number of scene properties (reflectance, motion, surface pattern, shape, pose, and lighting), each taking only a small number of values (e.g., completely matte or completely reflective). The value of this artificial world is complete access to the ground truth, both during the model’s training diet and when probing it subsequently. We do not intend the model as a general-purpose material recognition tool, and in fact find that it generalises quite poorly even to artificial stimuli outside its training distribution. Learning the far more varied scene properties that give rise to the far more complex temporal and spatial structure of the real world may only be a matter of scale, though, given the impressive abilities of commercial video prediction models (Wiedemer et al., 2025). Recent work shows that video-prediction models trained on real-world footage best capture shape-from-motion representations in the dorsal stream (Bai et al., 2026).

Our work focuses on whether learning by prediction can in principle discover untangled representations of material and other scene properties. The objective of predictive learning may be implemented in many different recurrent deep neural network architectures. We selected PredRNN (Wang et al., 2017) primarily for its video prediction quality, and do not intend its explicit splitting of units into separate temporal and spatiotemporal pathways as a hypothesis about the architecture of the visual cortex. Likewise, the predictive learning objective may be formulated in many different ways, and we are not committed to the prediction occurring at the lowest sensory level (in our model, pixels). Learning to predict in a latent space may be even more powerful (Bardes et al., 2024; Garrido et al., 2025). More generally, it seems likely that prediction is just one of several (or even many) learning objectives that structure internal representations, such as compression or efficient coding (Barlow, 1961; Olshausen & Field, 1996; Storrs et al., 2021), contrastive learning (Chen et al., 2020; Prince et al., 2024; Schneider et al., 2021; Sermanet et al., 2018), motion straightening (Niu et al., 2024), and others. Indeed, there is evidence that perceptual and neural responses are structured by both predictive (Lotter et al., 2020) and straightening (Hénaff et al., 2019, 2021) objectives.

Taken together, our results suggest that predictive learning discovers human-relevant material cues and provides a framework for understanding how brains learn rich scene representations without ground-truth world information. Learning to predict how visual experience changes over time is a feasible way for brains to discover underlying scene properties—like material—without access to ground-truth information.

## Methods

### Dataset

The dataset consisted of 20,000 twenty-frame greyscale videos of scenes procedurally generated in the Unity3D graphics engine (version 2019.4.27), each with a resolution of 128×128 pixels. Each scene contained one of 10 distinct objects, from meshes created in Blender (version 4.0), placed in front of a grey wall. Each scene was illuminated by one of 10 different lighting environments (4k HDR images downloaded from www.polyhaven.com), with a weak spotlight as an additional light source to cast an object shadow at a random angle. The albedo of the back wall was randomly sampled to vary the contrast and polarity of the object against its background. Objects either had a fully reflective surface (50% probability) or a matte surface (50% probability) textured with one of 10 different patterns (Figure 2). Objects were positioned close to the virtual camera at a slightly jittered distance and a random initial pose, and were either translated by a random distance (up to half the width of the image) or rotated by a random angle (up to 180 degrees) along all three axes over the course of 20 frames. This created close-up video clips of curved surfaces, with either matte or specular optic flow, moving in diverse ways in diverse lighting conditions (see Supplementary Videos for example training videos).

### Training and Testing the Model

We trained a four-layer recurrent network to predict the pixels of the next frame in each video. Specifically, we trained a PredRNN (Wang et al., 2017) recurrent convolutional network (Figure 3a) with 128 feature maps in each layer to minimise mean squared pixel error between predicted and actual future frames. We used the first 17,250 videos to train the network. For frames 1-10, the network predicted one frame into the future; after frame 10, it predicted until frame 20 with no further input. Each frame was resized into non-overlapping 4 ✕ 4 patches, which resulted in a 32 ✕ 32 spatial grid with 16 channels before passing through the network. During training, we used reverse scheduled sampling to control whether the network utilizes the ground-truth frames or generated frames at different time points. The reverse scheduled sampling started changing from the 5,000th step until the 50,000th step with α = 2,000, creating a gradual transition from 50% ground-truth frames early in training to 100% late in training. The model was optimized using the Adam optimizer with a learning rate of 0.0001 and a batch size of 12. The model was additionally trained on reversed frame sequences to augment the training.

We evaluated the model and extracted latent activations for analysis using the remaining unseen twenty-frame video sequences. As during training, the first 10 frames at test time were driven by real inputs, and frames 10-19 were driven by the model’s previously generated frames. Thus, the early steps reveal the processing of real input, and the later steps reflect the model’s longer-term predictions. We saved the latents from all three memory types (C-state, H-state, and M-state) in zero-indexed layers 1, 2, and 3, which correspond to the second, third, and fourth recurrent layers in the network. C-state corresponds to the temporal cell state of the network that holds short-term information which flows only “horizontally” to the same layer at the next time step. H-state is called the hidden state, and it is responsible for longer-term memory retention that can propagate both vertically across layers and horizontally across time. M-state is the unique spatiotemporal memory state introduced by PredRNN (Wang et al., 2017). It captures spatiotemporal details and can propagate this information in a zigzag format across layers and memory states to create a better spatial representation. We saved activations for frames 1-19 for each video, with each layer involving 128 feature maps that have spatial dimensions of 32 ✕ 32. Therefore, the resulting activation data for one video had dimensions of 19 ✕ 128 ✕ 32 ✕ 32 for each layer and memory type.

### Diagnostic Dataset

To test whether the model encodes material like a human, we created a new diagnostic dataset. By using the same poses, illuminations, and objects in Unity3D, we created 90 matched sets of three twenty-frame videos. In the first version, the object has a shiny mirror-like surface, which reflects different parts of the environment as the object rotates along a random axis. In the second version, the same object rotates along the same axis, but the reflections are rendered “stuck” to the object, moving with the surface like texture markings. The 10th frame of the “mirror” and “sticky” versions is identical. This shared frame is duplicated to create the third version, a stationary condition in which the object is static throughout. These matched sequences were rendered by rotating the camera and rear wall around a static object, rather than moving the object. This created the normal appearance of a moving shiny object in the first condition. In the second, “sticky reflections,” condition, it allowed us to render the reflections relative to a virtual viewer, who stayed static along with the object, while the camera moved around it. This causes the reflections to stay static relative to the object, which creates the impression of markings “painted onto” a matte surface (Supplementary Videos).

### Data Analysis

#### Evaluation of Prediction Performance

The video prediction performance of the network was calculated on 2,750 sequences, each consisting of 20 frames. For each sequence, the first ten frames were provided to the network, and then the subsequent frames were predicted autoregressively. The prediction performance was defined as mean squared error (MSE), which was calculated as the average pixel-wise squared difference between the luminance level in the predicted and the ground-truth frame. Pixel intensities were normalized to [0,1].

We compared the network’s performance against a persistence baseline and an optical flow baseline (RAFT). The persistence baseline was calculated by using the last observed frame (10th frame) to predict the next frame (11th frame). Each subsequent frame was then predicted from the previously predicted image, without access to the ground-truth frame. For the optical-flow baseline, a pretrained RAFT model (Teed & Deng, 2020) was used, enabling us to estimate optical flow between the last two observed frames (9th and 10th frames). This estimation was then used to predict the 11th frame by warping the 10th frame. For the following extrapolations, the previously predicted image was warped using the same flow field, without providing ground-truth images. Image warping was applied using bilinear interpolation.

#### Extraction of activations within the neural networks

Recurrent state activations were saved for 9,500 sequences from 10,500-19,999 of the dataset. The latent activations included 6,750 training sequences, as well as 2,750 sequences that were not used during the PredRNN training. For decoding and lesioning analyses, classifiers were trained on only the latents corresponding to sequences used during PredRNN training. The PredRNN-unseen subset of the latents was used to evaluate the accuracy of decoding and the impact of lesioning. The full latent set was used for tSNE visualizations.

Static scenes were not used for the decoding and lesioning analyses, as they served to increase variety during training and were not representative of the full dataset (10% of 20,000 videos). Objects with shiny surfaces were excluded from the decoding and lesioning analyses for the “surface pattern” variable, as their mirror-like material reflects the environment and has no defined surface pattern. Likewise, matte objects were excluded from analyses of the “illumination” variable since, without reflections, they lacked visual information about the lighting environment. Finally, the separate analysis of each layer and memory type did not reveal substantial differences; therefore, we combined all layers and memory types for all analyses except for single-unit decoding.

#### Visualization of Activations using t-SNE

We utilised t-SNE to visualize how the network represents and clusters object materials and motion characteristics at certain time points. The raw activations were spatially averaged across each 32 ✕ 32 convolutional feature map and then normalized across video sequences, before being provided to the tSNE algorithm. We implemented it in scikit-learn (Python) to embed the activation into two dimensions, with a fixed random seed (random_state = 0) and all other parameters set to default.

#### Linear Decoding Analyses

Linear decoding analyses were performed to examine the network’s performance in distinguishing the conditions of each scene variable: motion (rotating or translating), material (shiny or matte), shape (which of 10 objects), illumination (for shiny objects: which of 10 lighting environments), and surface pattern (for matte objects: which of 10 surface textures). Linear classifiers were implemented using logistic regression (scikit-learn, Python), with L2-regularisation, balanced class weights, and a maximum of 2,000 iterations. Features were standardised by the training-set mean and standard deviation. The same transformation was applied to the test data. No cross-validation was performed. Classifiers were trained and evaluated on different training and test sequences.

For the population decoding, raw latent activations were averaged across each 32 ✕ 32 convolutional feature map, then concatenated across the three layers and memory types. For each scene variable, a separate classifier was fit to the representations from each frame of the training sequences and tested on the corresponding frame of the test sequences. The population decoding performance was quantified by the area under the receiver operating characteristic curve (AUC) for each scene variable per frame, revealing the temporal dynamics of the network’s decoding performance. For single-unit decoding, raw activations from frame 10 were averaged across each 32 ✕ 32 convolutional feature map, but layer and memory type were preserved. For each scene variable, 1,152 separate classifiers were therefore fitted (3 layers ✕ 3 memory types ✕ 128 features) using unseen test sequences. For binary scene variables (motion and material), AUC values were used to define performance, with selectivity in either direction characterised by the absolute deviation from chance (0.5). For multiclass variables, the maximum one-vs-rest AUC across classes was used. Statistical significance was measured by 1,000 label permutations per unit, with a Benjamini-Hochberg false discovery rate correction. The data from all layers and memory types are shown combined, since there was no noticeable difference between layers or memory types.

### Feature Map Lesioning Analysis

To perform feature-specific lesioning analyses, we fitted separate linear decoders to the spatially averaged activations from frame 10, concatenated across all three layers and memory types (1,152 feature maps). These linear decoders were trained on the representations of training data and evaluated on the unseen test sequences. For each scene variable, the magnitude of weights of each feature map was ranked from highest to lowest. Then, we selected the feature maps with the top 10% of the highest weights to ablate, which corresponds to the most relevant feature maps for the scene variable under investigation. These feature maps were replaced with the mean activation from the training data, which amounts to zero after standardization. We then repeated this analysis, this time lesioning not the highest-weighted units, but 10% of units randomly selected from the whole population, over 500 repetitions. The random lesioning acts as a control condition, allowing us to quantify how large the performance drop caused by targeted lesioning is, compared to that expected when losing a matched number of non-targeted units. The impact of lesioning was calculated as the proportional drop in AUC relative to the baseline. To do that, we calculated AUC values before lesioning, after targeted lesioning, and after random lesioning. We retained the original weights throughout the analysis without refitting the classifiers after lesioning, which allowed us to assess how much the decoder’s performance depends on the selected feature maps. Statistical significance was measured by one-sided empirical randomization tests, with Holm correction applied across five scene variable comparisons.

We also wanted to test whether the most informative feature maps for different scene variables share overlapping representations. To do so, we selected the top 10% of feature maps (116 out of 1,152) based on the analysis described above. For each pair of scene variables, we computed the percentage of shared feature maps. To analyse whether the observed overlaps were greater than chance-level, we used one-sided exact stratified randomization tests with Holm-Bonferroni correction (α = 0.05), allowing us to preserve the exact number of selected feature maps across all memory types and layers. The hypergeometric distributions were used to calculate the null distributions.

## Data and code availability

All data from human observers and model predictions will be made public on GitHub and Zenodo, along with code to implement the models/weights of the pretrained network

## Acknowledgements

This research was supported by the Royal Society of New Zealand (Marsden Fast Start grant MFP-UOA2109 to K.R.S), and Germany’s Excellence Strategy via the Deutsche Forschungsgemeinschaft (EXC 3066/1 “The Adaptive Mind” – Project No. 533717223), as well as by the European Research Council (ERC) Advanced Award ‘STUFF’ (project number ERC-ADG-2022-101098225 to R.W.F.). The funders had no role in study design, data collection and analysis, decision to publish, or preparation of the manuscript.

## Author contributions

**F.K.:** Conceptualization, Methodology, Analysis, Writing—Original Draft. **R.W.F.:** Conceptualization, Writing–Review & Editing, Supervision, Funding acquisition. **K.R.S.:** Conceptualization, Methodology, Software, Analysis, Investigation, Writing–Original Draft, Visualisation, Funding acquisition;

